# SARS-CoV-2 variants exhibit increased kinetic stability of open spike conformations as an evolutionary strategy

**DOI:** 10.1101/2021.10.11.463956

**Authors:** Ziwei Yang, Yang Han, Shilei Ding, Andrés Finzi, Walther Mothes, Maolin Lu

**Author notes:** Correspondence to: Maolin Lu. These authors contributed equally to this work (Ziwei Yang, Yang Han).

## Abstract

SARS-CoV-2 variants of concern harbor mutations in the Spike (S) glycoprotein that confer more efficient transmission and dampen the efficacy of COVID-19 vaccines and antibody therapies. S mediates virus entry and is the primary target for antibody responses. Structural studies of soluble S variants have revealed an increased propensity towards conformations accessible to receptor human Angiotensin-Converting Enzyme 2 (hACE2). However, real-time observations of conformational dynamics that govern the structural equilibriums of the S variants have been lacking. Here, we report single-molecule Förster Resonance Energy Transfer (smFRET) studies of S variants containing critical mutations, including D614G and E484K, in the context of virus particles. Investigated variants predominantly occupied more open hACE2-accessible conformations, agreeing with previous structures of soluble trimers. Additionally, these S variants exhibited decelerated transitions in hACE2-accessible/bound states. Our finding of increased S kinetic stability in the open conformation provides a new perspective on SARS-CoV-2 adaptation to the human population.

## INTRODUCTION

Since the beginning of the severe acute respiratory syndrome coronavirus 2 (SARS-CoV-2) pandemic, variants of concern (VOCs) that are more transmissible and possibly more pathogenic have emerged and caused devastating consequences worldwide. The currently administrated interventions, vaccines and antibody therapy are directed against the virus surface spike (S) protein as the primary target of antibody response ^1-6^. However, emerged and emerging SARS-CoV-2 variants harbor accumulated mutations in S (Figure 1), including the D614G (S_G614_), U.K. B.1.1.7 (S_Alpha_), South African B.1.351 (S_Beta_), Brazilian P.1 (S_Gamma_), and Indian B.1.617.2 variant (S_Delta_), of which some have gained increased resistance to antibody neutralization ^7-16^. While current vaccines remain efficient against severe disease induced by these VOCs ^17,18^, additional mutation in S could compromise this protection. Similarly, some VOCs are now resistant to first generation antibody therapies and impacted several molecular tests ^11,12,19-21^. For instance, D614G is a spike variant that first emerged in January 2020 and became a dominant form as the pandemic spread, reaching >74% global prevalence by June 2020. D614G has been associated with increased infectivity and transmissibility of SARS-CoV-2 ^22,23^. E484K has been identified as an escape mutation that facilitates virus immune evasion ^24^. E484K was first identified in South Africa (B.1.351) and Brazil (P.1), but it was also gradually adopted by the U.K. (B.1.1.7) strain since, indicating an associated selective advantage ^23^. Other mutations on S variants such as L452R, N501Y, and P681R also attracted research attention due to enhanced ACE2 interaction and partial escape from vaccine-elicited antibodies ^25^. Therefore, it is of great importance that we understand how these mutations affect the conformational landscape of S, and how this connects to the enhanced immune evasion, infectivity, and transmissibility of SARS-CoV-2 variants.

**Figure 1.**
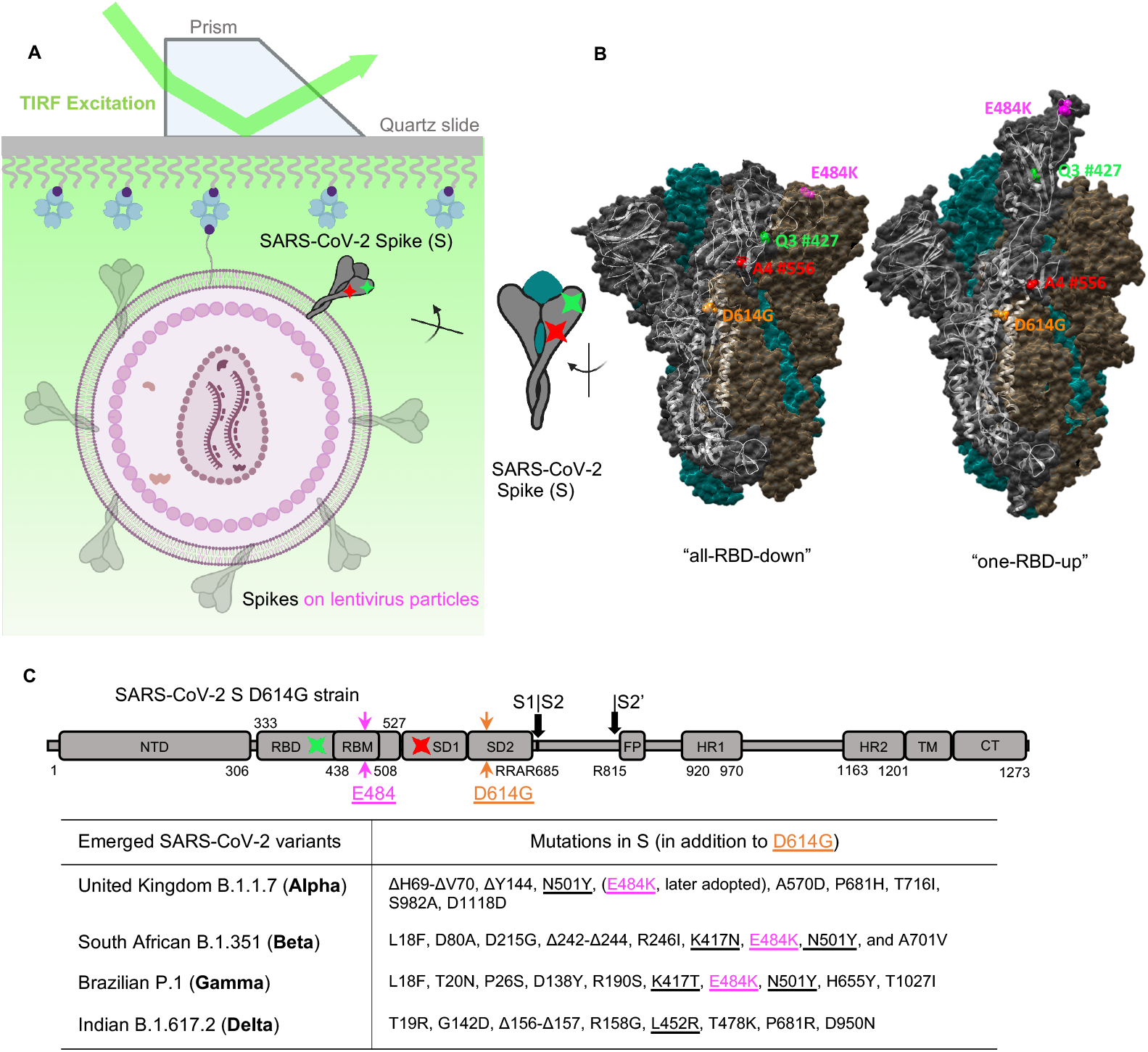
Experimental design for characterizing the conformational dynamics of the SARS-CoV-2 spike variants on virus particles via single-molecule FRET. (**A, B**) Experimental set-up. **(A)** Virus-like particles that carry a single two-dye-labeled SARS-CoV-2 spike protomer among unlabeled wildtype spikes were immobilized on a quartz slide and then imaged on a prism-based total internal reflection fluorescence (TIRF) microscope. The quartz slide was passivated with PEG/PEG-biotin to enable streptavidin coating and the subsequent immobilization of virions that contain a biotin-lipid (DSPE-PEG-biotin). The virus-like particles were composed of HIV-1 cores and SARS-CoV-2 spikes on the virus surface. **(B)** Positioning of labeling dyes for the site-directed incorporation of fluorophores (LD555-cadaverine: a Cy3B derivative, green; LD655-CoA: a Cy5 derivative, red) was elucidated by the S1 conformational change from the ‘‘RBD-down’’ to the ‘‘RBD-up’’ structures, which are adapted from RCSB PDB: 6ZB5 (‘‘all-RBD-down’’) and 7KJ5 (‘‘one-RBD-up’’). **(C)** Domain organization of the parental full-length SARS-CoV-2 spike protein with D614G and E484K mutation. The introduction sites of labeling tags Q3 (green) and A4 (red), where the donor and acceptor fluorophores will be conjugated to respectively, are indicated. SD1, subunit domain 1; SD2, subunit domain 2; S1/S2, S2’, protease cleavage sites; FP, fusion peptide; HR1/HR2, heptad repeat 1/heptad repeat 2; TM, transmembrane domain; CT, cytoplasmic tail. Labeling peptides Q3 and A4 are inserted into RBD and SD1 respectively before and after RBM within the S protein in these studies. Emerged variants that are currently classified by CDC as “variants of concern” and their respective key mutations in the spike are specified in the table.

The S protein is a trimer of S1/S2 heterodimers ^26,27^. The surface-exposing S1 contains the receptor-binding domain (RBD) that engages with cellular receptor human angiotensin-converting enzyme 2 (hACE2) ^28-31^, an event which, upon occurrence, triggers the transmembrane S2 to mediate the virus-cell membrane fusion process that enables virus entry ^30-34^. The S glycoprotein is structurally flexible and conformationally dynamic to facilitate both virus entry and antibody evasion. Each spike protomer samples two well-characterized RBD-orientation-based conformations, the hACE2-accessible “RBD-up” and the hACE2-inaccessible “RBD-down” conformations (Figure 1). Consequently, the S trimer can exist in an equilibrium of four possible combinations of the protomers - four conformational states of S trimer: “one-RBD-up,” “two-RBD-up,” “all-RBD-up,” and “all-RBD-down.” Extensive structural studies of truncated soluble and virus-associated parental/original strain S_D614_ have confirmed the existence of the above-mentioned major S trimer configurations ^29-31,35-43^. As S continuously evolves, accelerated efforts have been implemented to characterize emerging S variants genetically, virologically, and structurally ^25,44-54^. However, real-time monitoring of structural/conformational dynamics of S variants that connect structures has been lacking. The mechanism by which spike proteins adapt their conformations and dynamics during virus evolution could facilitate the understanding of pathogenesis, antibody neutralization, and vaccine efficacy of these S variants.

Using single-molecule Förster Resonance Energy Transfer (smFRET), we have experimentally revealed that virus-associated S_D614_ dynamically samples at least four distinct conformational states, transitioning in time from milliseconds to seconds ^55^. On the faster dynamic scale, a remarkably long ∼ 130 μs sampling simulation of the S_D614_ opening process suggested the existence of five conformations between the transition from “RBD-down” to “RBD-up,”^56^ despite the temporal limitation that existed in simulation. Nevertheless, real-time experimental observations of conformational landscapes and the dynamics that govern the conformational equilibriums of recently emerged S variants on the viruses have not been performed. It is unclear whether arising S variants differ from the original S_D614_ on viruses regarding conformational dynamics/kinetics. Here, we further deployed our smFRET approach to reveal the dynamic details and the mechanistic of the conformational shifts in different S variants at the surface of lentivirus particles. Spike variants that contain our mutations of interest, D614G or/and E484K, as well as their parental strains were analyzed and compared side-by-side in this study. Using smFRET, we investigated how these mutations affect the conformational landscape and dynamics of the S glycoprotein by characterizing the overall movements within S in the millisecond to the second range. We discuss how such influences contribute to the selective advantage that variants harboring these mutations confer, and provide insights into the mechanisms by which mutations in S could increase the severity of SARS-CoV-2 variants.

## RESULTS

### Establishing real-time monitoring of conformational profile of SARS-CoV-2 Spike (S) variants via smFRET

To perform smFRET investigations of spike variants, lentivirus particles that carry only a single FRET-paired dyes-labeled SARS-CoV-2 Spike protomer among unlabeled wildtype Spikes were imaged on a prism-based total internal reflection fluorescence TIRF microscope (Figure 1). Lentivirus particles contain the HIV-1 core and the SARS-CoV-2 S glycoproteins on the virus surface. As previously described ^55^, Q3 and A4 peptide tags were inserted before and after the receptor-binding motif of tested S variants at positions 427 and 556 (427-Q3/556-A4, Figure 1) to allow the site-specific introduction of donor and acceptor dyes/fluorophores onto S1 (see METHODS). We previously validated that Q3 and A4 introduced at the specified locations didn’t affect viral infectivity ^55^. Lentivirus particles used for smFRET imaging were prepared by transfecting 293T cells with a 20-fold excess of plasmid encoding S_D614_, S_G614_, S_Alpha_, or S_Alpha+E484K_ over their corresponding 427-Q3/556-A4 plasmid. Statistically, this strategy similarly used for the analogous investigations of HIV-1 envelope ^57,58^ ensures the production of virus particles that contain, on average, only a single Q3/A4-tagged FRET-engineered S protomer within a hybrid trimer (tagged - wildtype - wildtype) among elsewhere wildtype S trimers on the virus. Enzymes ^59,60^ that site-specifically recognize amino acids in respective Q3 and A4 tags transfer customized dye-conjugated substrates to these tags ^55^, so that the donor and acceptor dyes (LD555, green; LD655, red) were introduced into the S1 of S variants (Figure 1A). Fluorescently labeled lentivirus particles were then immobilized on a quartz slide and imaged on a prism-based TIRF microscope. Donor fluorophores were directly excited by a single-frequency (532 nm) continuous-wave laser, and fluorescence emissions from both donor and acceptor were separately recorded at 25 Hz (Figures 2B, 3C,E, and S1A-B). Conformational motions in the Spike will lead to changes in the inter-dye distance exampled by structural switching from “RBD-down” to “RBD-up” (Figure 1B), causing the donor-to-acceptor energy transfer efficiency (FRET efficiency) changes. FRET values can be derived by fluorescence signals of FRET-paired dyes in real-time. In smFRET experiments. We extracted fluorescence traces from the recorded movies that demonstrate anti-correlated levels of donor and acceptor fluorescence and derived FRET traces (Figures 2B, 3C, E, and S1A-B) to reflect conformational changes in the Spike structures.

**Figure 2.**
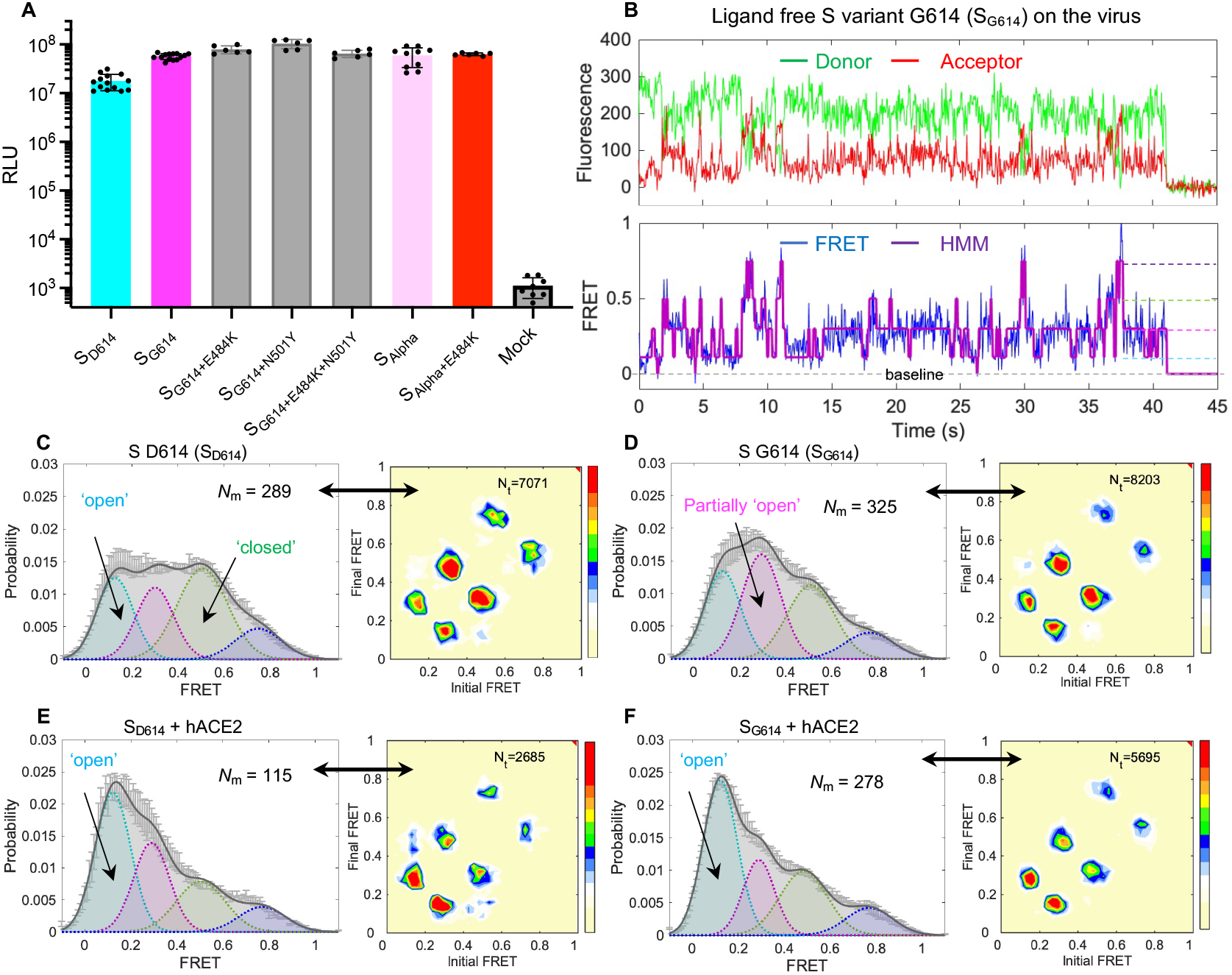
D614G substitution shifts the conformational landscape of unbound spike from the ground state to asymmetrically configurated intermediate states. **(A)** Infectivity quantification for HIV-1 lentivirus particles carrying various spike variants determined on hACE2 expressing 293T cells (293T-ACE2). Infectivity (mean ± s.d.) was measured from three independent experiments in triplicates. RLU, relative light units. **(B)** Example fluorescence trace (LD555, green; LD655, red) and resulting quantified FRET traces (FRET efficiency, blue; hidden Markov model initialization, red) of a dually labeled ligand-free spike protein on the surface of HIV-1 lentivirus particle. The single-step photobleaching step of dyes at the single-molecule level define the baseline (dashed black). Four distinguishable FRET-populated states are indicated as color-coded dash lines. (**C, D**) FRET histograms (left) and TDPs (right) of ligand-free D614 spike (S_D614_,**C**) and G614 spike (S_G614_, **D**) on lentivirus particles. A number (*N*_m_) of individual active/dynamic molecules - FRET traces were compiled into a conformation-population FRET histogram (gray lines) and fitted into a 4-state Gaussian distribution (solid black) centered at 0.1-FRET (dashed cyan), 0.3-FRET (dashed red), 0.5-FRET (dashed green), and 0.75-FRET (dashed magenta). TDPs show the distributions of initial and final FRET values for every observed transition in FRET traces. TDPs are displayed as initial FRET vs. final FRET with relative frequencies (max red scale = 0.01 transitions/second), originated from the idealization of individual FRET traces in FRET histograms. TDPs trace the locations of state-to-state transitions and their relative frequencies. (**E, F**) Experiments as in (C, D), respectively, conducted in the presence of 200 μg/mL monomeric hACE2. The soluble hACE2 activates spike proteins on the virus by shaping the conformational landscape toward stabilizing the “all-RBD-up” conformation (activated state). FRET histograms represent mean ± SEM, determined from three randomly assigned populations of FRET traces under corresponding experimental conditions. *N*_m_, number of individual FRET traces. Evaluated relative state occupancies see Table S1.

### D614G substitution induces a conformational shift of virus-associated spikes towards receptor-accessible states

We first tested S-dependent infectivity of lentivirus particles on hACE2 expressing 293T cells (293T-ACE2). Virus particles bearing S_G614_, S_G614+N501Y_, S_G614+E484K_, S_G614+E484K+N501Y_, S_Alpha_, or E484K carrying S_Alpha_ (S_Alpha+E484K_) variants have a discernible increase in infectivity compared to viruses bearing the parental S_D614_ (Figure 2A) in agreement with previous results ^49,50,53,61,62^.

To know whether D614G substitution on S would influence the conformational profiles of Spikes, we next performed smFRET analyses of S_G614_ on lentivirus particles compared to the original S_D614_. The example fluorescence donor/acceptor trace and the derived FRET efficiency trace of an individual S_G614_ on the virus (Figure 2B) shows that S_G614_ primarily occupies the 0.3-FRET (dashed red line)-indicated conformation and constantly interconverts between four conformational sates indicated by four different FRET levels (01-FRET, 0.3-FRET, 0,5-FRET, and 0.75-FRET). We have previously shown that S_D614_ on both lentivirus-like and coronavirus-like particles exist in four distinct FRET states: the 0.5-FRET state is the mostly abundantly occupied and it corresponds to the “all-RBD-down” closed conformation; the 0.3-FRET state corresponds to the “one/two-RBD-up” partially open fusion-promoting conformation; the 0.1-FRET state corresponds to the “all-RBD-up” fully open fusion-activated conformation; and the high-FRET (slightly varied values) is an unknown state ^55^. We confirmed our reported results of S_D614_ primarily being in the “all-RBD-down” conformation from newly-generated conformational population-indicated FRET histogram (Figure 2C), compiled of hundreds of FRET traces. In contrast, the FRET histogram of unbound S_G614_ on lentiviral particles demonstrate that S_G614_ inherits all four conformations from the S_D614_ with a shift in the conformational landscape (Figure 2D). S_G614_ exhibits a propensity to occupy the 0.3-FRET indicated “one/two-RBD-up” partially open states, in which either one or two RBD is exposed for hACE2-binding (Figure 2D). Thus, D614G substitution shifts the conformational landscape of the original S_D614_ from hACE2-inaccessible “all-RBD-down” conformation dominance to predominately hACE2-accessible conformations. Our findings are consistent with the observations of increased occupancy of “RBD-up” structures characterized by cryoEM, despite the variations in quantification among all published structural work ^48-51,61^. In the presence of the hACE2, both S_D614_ and S_G614_ adopt the 0.1-FRET dominant histograms (Figures 2E and F), meaning that higher proportion of Spikes on the virus surface are in “all-RBD-up” fully open fusion-active conformations. The relative state occupancies were quantified in Table S1. The propensity towards the fully open conformation (0.1-FRET) can be directly observed from the individual FRET traces (Figure S1).

Next, we wanted to gain insights into the sequence and the frequency of conformational dynamics of S_D614G_. We applied the Hidden Markov Modeling (HMM) ^63^ to idealize all FRET traces compiled into the FRET histogram individually and quantitatively analyzed thousands of transitions observed for the S glycoproteins on the viruses. The order and the frequency of transitions were displayed on transition density plots (TDPs), as distributions of initial FRET vs. final FRET of all transitions (Figures 2C-F, right panels). The comparison of TDPs between S_D614_ and S_G614_ (Figures 2C-D, right panels) show that conformational changes for S glycoproteins exhibit a defined sequence from the “all-RBD-down” closed state (0.5-FRET) to the “all-RBD-up” activated state (0.1-FRET) through a necessary intermediate 0.3-FRET state – “one/two-RBD-up”. This finding implies that the D614G substitution does not alter the sequence of events that constitute the conformational dynamics of the spike, although it does induce shifting of the conformational landscape/distribution (Figures 2C and D). Similarly, the sequence of conformational transitions of hACE2-bound S-D614 or S-G614 remains identical to the ligand-free Spikes (Figures 2 and 3). The conformational distribution of the Spike population, however, shifts towards an “all-RBD-up”-predominant state upon hACE2-binding (Figures 2E and F).

**Figure 3.**
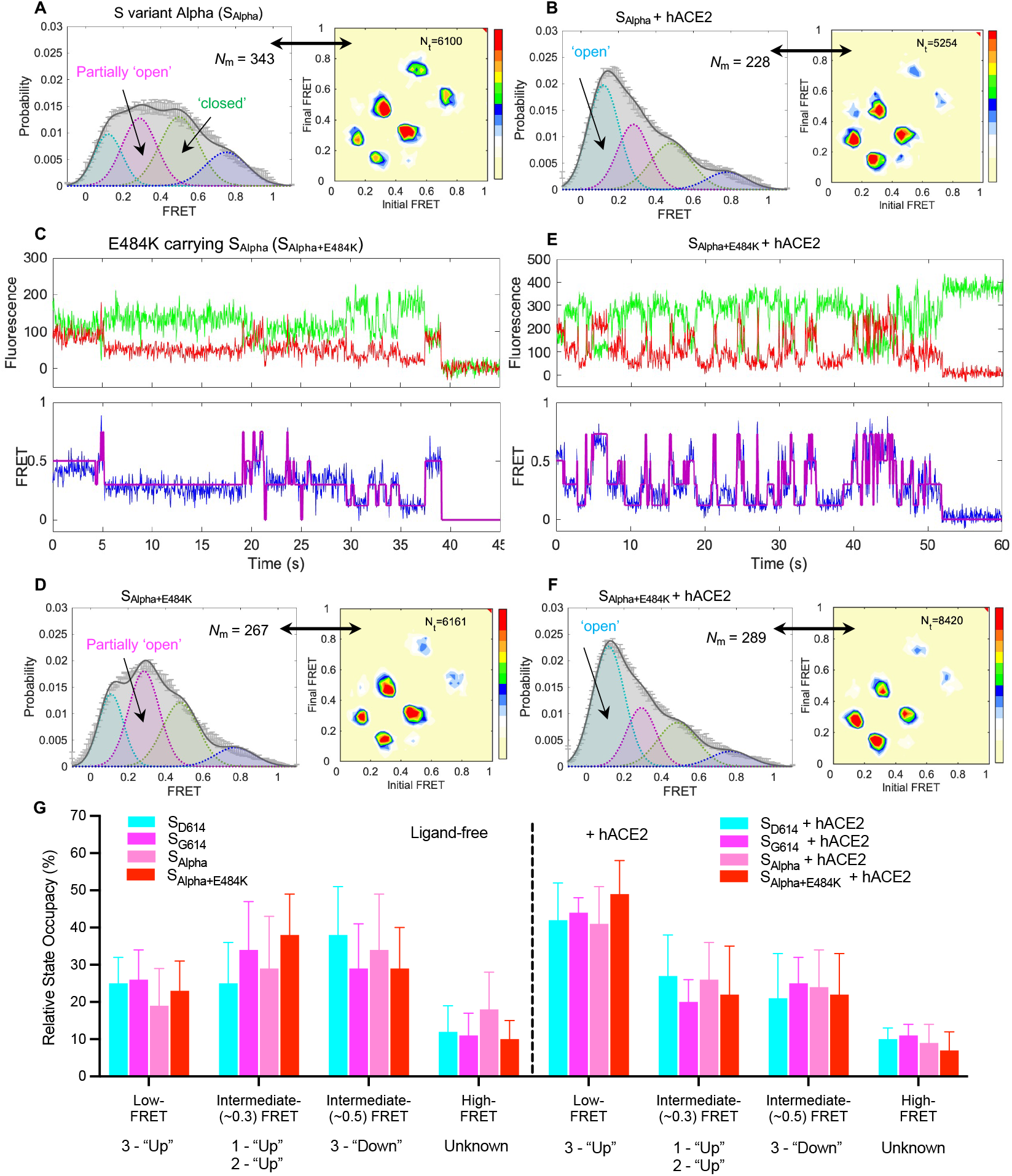
E484K stabilizes the S Alpha variant towards ‘RBD-up’ conformations. (**A, B**) FRET histograms (left) and TDPs (right) of S Alpha variant (S_Alpha_) on lentivirus particles with (**A**) and without (**B**) 200 μg/mL hACE2 presence. The soluble hACE2 activates the S_Alpha_ by shaping the conformational landscape toward stabilizing the “all-RBD-up” conformation. **(C)** Representative fluorescence traces (LD555, green; LD655, red) and quantified FRET traces of a single ligand-free E484K carrying S Alpha variant (S_Alpha+E484K_) on lentivirus particles. **(D)** The FRET histogram (left) and TDP (right) of ligand-free S_Alpha+E484K_ on lentivirus particles. (**E, F**) Experiments as in (C, D), conducted in the presence of 200 μg/mL hACE2. FRET histograms represent mean ± SEM, determined from three randomly assigned populations of all FRET traces under corresponding experimental conditions. *N*_m_, number of individual FRET traces. (**G**) Quantification of the FRET-indicated state occupancy for different spike variants. The occupancy in each FRET state was presented as mean ± SEM, determined by estimating the area under each Gaussian curve in FRET histograms. Fitting parameters see Table S1.

### E484K carrying S_Alpha_ resides predominately in partially open “one/two-RBD-up” receptor-accessible conformations

E484K and N501Y reside in the receptor-binding motif (RBM) of Spike that directly contacts the receptor hACE2 (Figure 1B). All three S variants, S_Alpha_ (later strain), S_Beta_, and S_Gamma_ share both N501Y and E484K substitutions (Figure 1C), in which E484K was later adopted by Alpha variant after the emergence of Beta and Gamma variants. Increasing experimental evidence indicates that E484K enables immune evasion, while N501Y may foster increased virus transmissibility by enhancing hACE2 binding ^7,8,24,44,64,65^. Analysis of FRET histogram and TDP of ligand-free S_Alpha_ on lentivirus particles revealed that S_Alpha_ (Figure 3A) sampled slightly more “one/two-RBD-up” (partially open conformations) on the viruses than the S_D614_ but less than S_G614_ (Figures 2C, 2D, and Table S1). This finding partially contradicts the cryoEM structural results ^52,53^, which indicate soluble forms of S_Alpha_ being more “open” compared to the same engineered soluble forms of S_D614_ and S_G614_. This insight has resulted from a lack of consideration of the activated state – a fully open “three-RBD-up” conformation. smFRET observed in situ conformations of S_Alpha_ incorporated on virus particles, whereas cryoEM characterized the soluble forms of S_Alpha_ protein. The differences in techniques likely contributed to the variations in the characterizations of the conformation-occupancy. Encouragingly, parallel in situ characterizations of two antibody-bound S_Alpha_ by smFRET and cryoET resulted in an overall agreement in the number of individual RBD units with different conformations ^54^, despite that smFRET is a dynamics-emphasizing approach whereas cryoET and other structural tools emphasize static features. Overall, the hACE2 binding resulted in shifting of the S-alpha conformational landscape towards a more fusion-active “all-RBD-up”-dominating state as expected (Figure 3B).

To further assess the conformational effects of E484K in the context of S_Beta_, S_Gamma_ and S_Alpha_, which later adopted the alternation, we next performed smFRET studies using E484K carrying S_Alpha+484K_ (Figures 3C-F). E484K substitution exhibits a noticeable effect of reducing the occupancy of ligand-free spike variant in the 0.5-FRET state and increasing the 0.3-FRET state occupancy (Figures 3C, D). It suggests that E484K stabilizes the S variants towards the “one/two-RBD-up” conformation by reducing the “all-RBD-down” conformation. The overall conformational landscape of ligand-free S_Alpha+E484K_ (Figure 3D) is similar to that of S_G614_ (Figure 2D), with slightly elevated occupancy in the “one/two-RBD-Up” conformations. The observation that E484K renders S variants adopting more “one/two-RBD-up” conformations agrees with all existing high-resolution cryoEM structural results of E484K carrying S_Beta_ and S_Gamma 52,53_. The similarity between overall conformational distributions (FRET histograms) of S_Alpha+E484K_ and S_G614_ is also in line with the structural and biochemical observations that highlighted the consistency between the structural and biochemical profiles of the E484K-containing S_Beta_ and S_G614_ variants ^53^. Interestingly, structural characterization of hACE2-bound S variants has been rarely reported. In our study, smFRET investigations of ligand-free and hACE2-bound S variants were done in parallel in a lentivirus context. Analysis of hACE2-bound S_Alpha+E484K_ (Figures 3E, F) and other variants in this study demonstrates that the parental Spike and variants are unambiguously predominated by the “all-RBD-up” (0.1-FRET) activated state upon binding to hACE2 receptors (Figure 3G). The quantifications of relative conformation-occupancy for all inspected S variants are summarized in Figure 3G, and the determining parameters are listed in Table S1.

### Spike variants show a propensity towards “RBD-up” conformations with decelerated transition dynamics than that of “RBD-down”

Our conformational results of all inspected S variants and their hACE2-bound state were further supported by the contour plots (maps), which display the FRET values across the entire 15-second duration of FRET detection (including those photobleached, zero-FRET values) in an unbiased fashion (Figure 4A). Contour plots demonstrate an increased number of the S_G614_ variant occupying partially open “one/two-RBD-up” intermediates (0.3-FRET range) compared to the original S_D614_, whose spikes are more enriched in the “all-RBD-down” (0.5-FRET range). Similarly, the E484K-containing S_Alpha_ variant also exhibits a higher frequency of spikes occupying the “one/two-RBD-up” intermediates (0.3-FRET) state than wildtype S_Alpha_. Upon hACE2 binding, all the inspected variants exhibit an increase in spike density towards the low FRET range. It indicates that a shift of the Spike conformational landscape from an “all-RBD-down” closed conformation or “one/two-RBD-up” partially open conformation to an “all-RBD-up” fully open fusion-activated conformation has taken place. Together, this ensembled representation of spike conformational landscape corroborates the conclusion deduced from the histogram built upon visually identified noise-avoiding traces and suggests that our real-time observation of conformation-population is robust despite the limited capacity of identifying potentially transiently (below milliseconds) sampled states.

**Figure 4.**
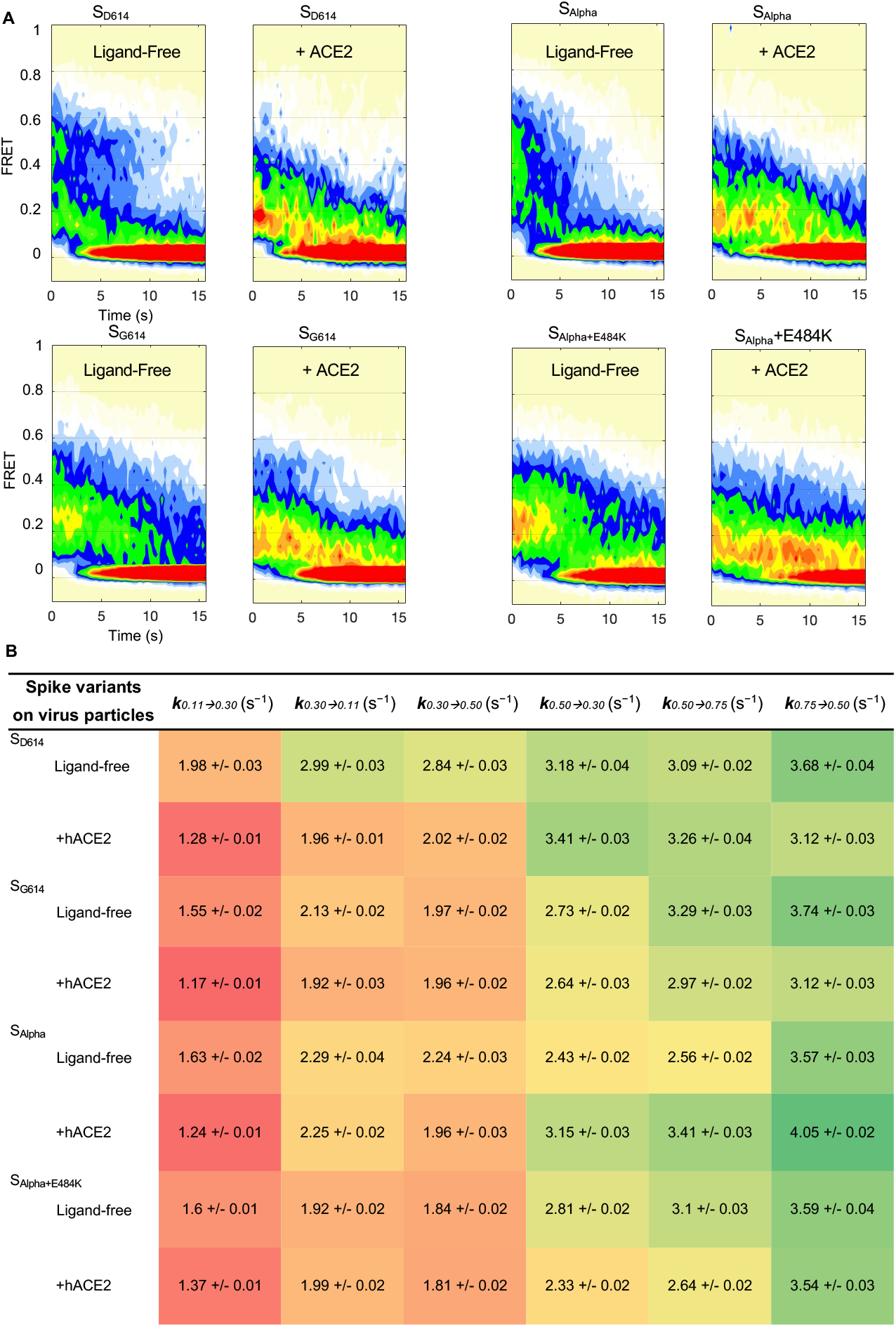
D614G and E484K render S on the virus in favor of ‘RBD-up’ open conformations, which are slower in temporal transitions. **(A)** Contour plots constructed from the compilation of the population of smFRET traces/trajectories. The contour plots were summed over time (15 seconds) of active/dynamic molecules compiled in the corresponding FRET histograms (Figures. 2C-F, 3A, B, D, and F). Some FRET-labeled active molecules that have reached photobleaching within 15 seconds contribute to the 0-FRET population. Contour plots blindly/unbiasedly monitor conformational ensembles occupied by virus-associated spike proteins over time. **(B)** Rates of transition between all observed FRET states for all tested S variants and their respective ligand-activated states. Rates are color-coded: warm-color indicates relatively faster, whereas cold-color means slower in transition. The distribution of dwell times (Figures S2 and S3) in each FRET state, determined through Hidden Markov Modeling (HMM), were fitted to the sum of two exponential distributions, *y(t)* = *A1* exp ^*–k1t*^ + *A2* exp ^*–k2t*^, where *y(t)* is the probability and *t* is the dwell time. The weighted average of the two rate constants from each fit is presented. Error bars represent 95% confidence intervals propagated from the kinetics analysis.

One of the unique features of smFRET analysis is to experimentally evaluate the transitioning kinetics among conformations of S variants in the context of virus particles by determining the transition rate between spike conformations (Figure 4B). The dwell times of each FRET-defined conformation describe the duration of time that each spike stays within a specific conformation before changing into the next. They were first identified by applying Hidden Markov Modeling (HMM) and were then compiled into dwell-time distributions. Dwell-time distributions were displayed in survival probability plots (Figures S2, S3) as the probability function of time-durations of spike occupying a particular conformation in a directional transition event. Two-component exponential decays were the simplest model to interpret our data. The weighted transition rate derived from the exponential fitting for each transition event signifies the speed at which the spike transitions from the current conformation to the next. The final rate constants for all transition events were listed in a heat map (Figure 4B), in which the smaller numbers are colored in red and larger numbers in green. The heat map of transition rates of different S variants amplifies subtle differences and reveals an exciting feature hidden in the interconverting map among four distinct conformations. We observed decreases in transition rates as the spikes progress sequential changes from the “all-RBD-down” closed state on the right to the “all-RBD-up” fully open activated state on the left (Figure 4B). It indicates that the spike trimer tends to dwell longer in hACE2-accessible “one/two/all-RBD-up” conformations. The finding of S_G614_ sampling more dynamics-reduced hACE2-accessible conformations implies its increased binding competence of spikes to hACE2 and enhanced kinetic stability compared to S_D614_. This likely explains the inconsistency reported previously surrounding hACE2-binding affinity and structural flexibility or stability of the soluble S_G614_^48-51,61^.

Our results of transition rates among four different conformations imply that the spike protomer is prone to be more conformationally stable in the “RBD-up” conformation than in the “RBD-down.” Thus, it is not surprising to observe that hACE2-bound spikes (“all-RBD-up”) show least propensity to shuttle back to the upstream conformations on the fusion path (Figure 4B). Similarly, the “all-RBD-up”, hACE2-bound conformation shows the lowest relative free energy in the derived quantitative model for hACE2 activation (Figure 5) from smFRET results. The ligand-free S_G614_ and S_Alpha+E484K_ spikes primarily occupy the “one/two-RBD-up” partially open state (0.3-FRET), which presents the lowest relative free energy among all four FRET-defined states. In contrast, all hACE2-bound spikes exhibit the lowest relative free energy at the 0.1-FRET state – the most populated “all-RBD-up” fully open fusion-activated state. Of note, our observation is solely on the S1 subunit and our results cannot inform the downstream conformations after the fusion-active open conformation of S1 with three RBDs oriented up. Taken together, smFRET analysis of S variants revealed that S in the “RBD-up” (hACE2-accessible) conformations/structures exhibit increased conformational stability with decelerated dynamics, providing a new perspective of molecular interpretations that S variants evolve to favor “RBD-up” conformations.

**Figure 5.**
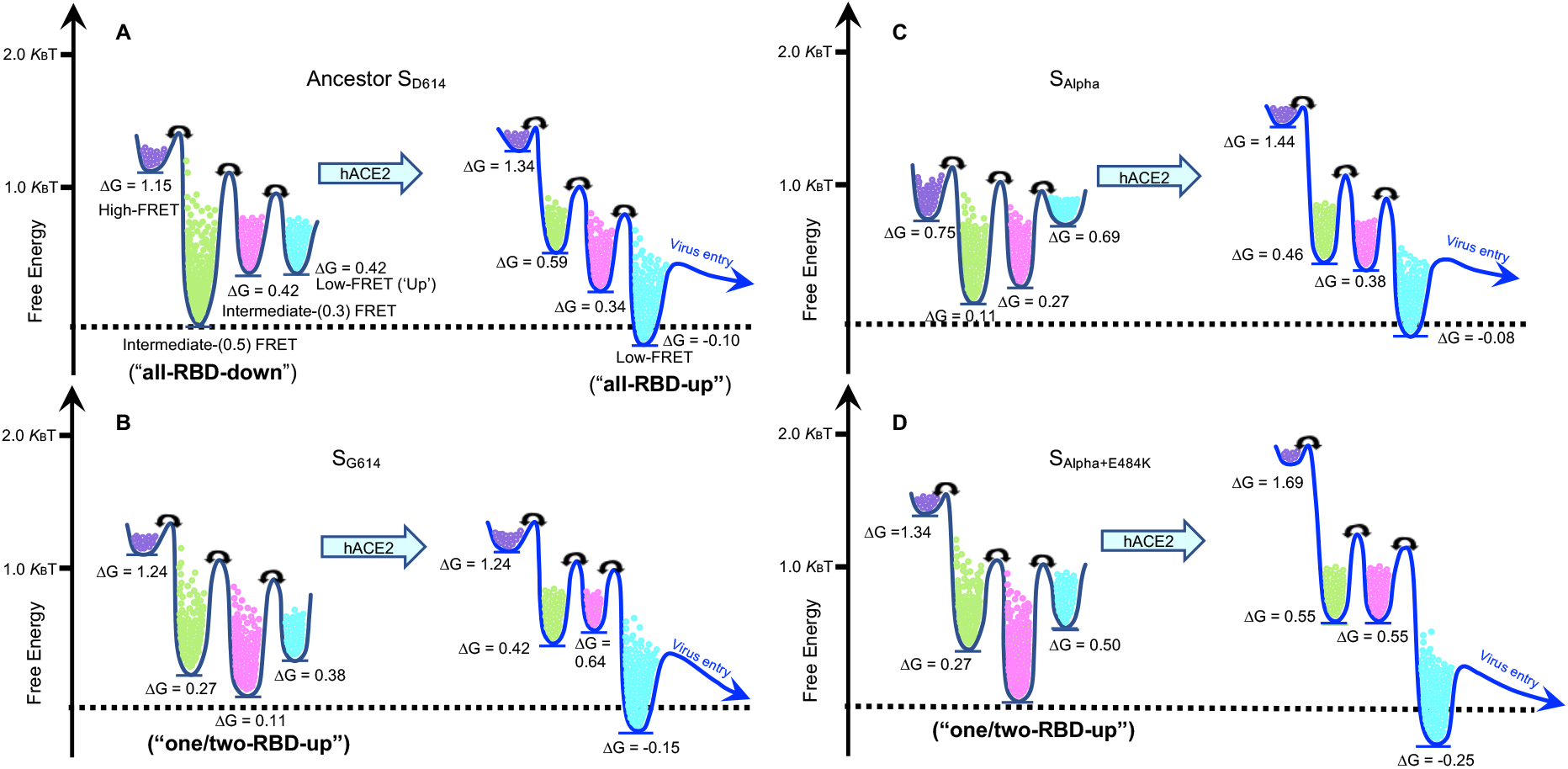
Relative free-energy models depict conformational landscapes of different virus-associated spike variants upon activation by the binding of hACE2. (**A - D**) Free-energy models of parental S_D614_ (**A**) and variants S_G614_ (**B**), S_Alpha_ (**C**), and S_Alpha+E484K_ (**D**). The differences in free energies between states were roughly scaled based upon the relative state occupancies of each state. The ligand-free D614G and E484K carrying spikes are dominated by the intermediate-FRET state (“one/two-RBD-up”), which exhibit the lowest relative free energy among all four FRET states. In contrast, all hACE2-bound spikes show the lowest relative free energy at the low-FRET state (“all-RBD-up”).

## DISCUSSION

SARS-CoV-2 variants of concern – VOCs (Figure 1C) pose significant challenge to public health worldwide. All four variants feature an initially world-wide adopted D614G mutation compared to the original Wuhan strain, while three of them contain an E484K mutation (E484K was also later adopted by the Alpha variant according to the World Health Organization, https://www.who.int/en/activities/tracking-SARS-CoV-2-variants/). As these variants have out-competed the original Wuhan strain and became the globally dominant strains, it is of great importance that we understand how these mutations within the spike influence the behavior and functionality of the spike. In this study, we have uncovered various aspects of such influence, which furthered our understanding of SARS-CoV-2’s survival strategy during the pandemic that started in 2019.

### The controversy of D614G on spike conformations and receptor-binding: enhancing conformational flexibility or stability? Lower or higher binding affinity to hACE2?

D614G mutation in the S protein is associated with enhanced transmissibility of SARS-Cov-2 viruses ^66,67^. We showed that lentiviruses that contain D614G spike exhibit significant increase in infectivity compare to the original Wuhan strain. High-resolution cryoEM structures of truncated soluble S_G614_ suggest that the naturally occurring S_G614_ is more conformationally flexible and favors the “open” conformations that are more accessible to hACE2, which likely accounts for enhanced virus transmissibility ^49-51,61^. A most recent cryoEM work shows that full-length S_G614_ exhibits higher S trimer stability and lower binding affinity to hACE2 than the parental D614 ^48^, which contradicts several studies ^49,50,67^. The coexistence of enhanced transmissibility and reduced hACE2 binding was rationalized by attributing the transmissibility primarily to the effect of D614G substitution on preventing S1 from shedding off on the virion, which consequently increases the number of functional spikes on the viral surface ^48^. This explanation was supported by observing a disordered-to-ordered transition of a “630 loop” caused by the D614G substitution, which seemingly stabilizes S trimer in the “one-RBD-up conformation”, preventing premature S1 shedding ^48^.

Using smFRET, we have determined that the S_G614_ variant exhibits a higher number of spikes residing in partially open conformations compared to S_D614_, consistent with all available structural data ^48-51^. Spikes sampling more partially open “one-RBD-up” or “two-RBD-up” hACE2-accessible conformations could potentially favor more hACE2 binding towards the fully fusion-activated “all-RBD-up” conformation ^49,50^. The observed decelerated transition rate of hACE2-accessible “RBD-up” conformations (Figure 4B) can contribute to S trimer stability ^48^. Thus, our results of S_G614_ dwelling longer within more hACE2-accessible conformations likely reconcile the controversy of hACE-binding and structural flexibility/stability of the soluble S_G614 ^48-51,61,67^_. The combined effect of D614G on enhancing S-hACE2 binding competence and stabilizing spikes in receptor-accessible conformations may confer the enhanced transmissibility of D614G bearing SARS-CoV-2 VOCs.

### Changing SARS-CoV-2 evolutionary adaptation: S-associated virus transmissibility vs. immune evasion

The original S strain adopted a predominately “all-RBD-down” closed conformation in order to conceal the RBD from host humoral responses ^29,31,42,55^, thus allowing the SARS-CoV-2 to better survive within the host. While S can only bind to hACE2 when the RBD is exposed, the concealment of RBD can be an important immune evasion strategy for the virus, as RBD is a primary target of neutralizing antibodies. As a cost of this protective trait, the receptor-binding ability of S has also suffered reduction due to the concealment of RBD. However, the subsequent variants carrying the D614G mutation within S have adopted another strategy to compromise this. D614G reduces S1 shedding and exhibits more functional spikes on the surface predominantly in hACE2-accessible conformations, which significantly increases the hACE2-binding competence, as discussed above. While this may contribute to an enhanced transmission rate, the increased number of surface S combined with their RBD-exposed conformation could render the viruses markedly more susceptible to host antibody attacks. It indicates that higher receptor-binding competence and faster transmission rate confer more fitness advantage for the spread of SARS-CoV-2 than a higher degree of protection against host immunity. Therefore, early SARS-CoV-2 evolution to the human host from late 2019 to early 2020 likely prioritizes receptor-binding competence to enhance transmissibility instead of the defense against host immunity.

In addition to D614G, E484K is another mutation adopted by most of the dominating strains, appearing in 3 of the 4 VOCs (Alpha, Beta and Gamma), of which Alpha variant later was similarly observed to adopt E484K according to the World Health Organization. The appearance of both E484K and N501Y mutations appear to increase the hACE2-binding affinity of S ^53,62^. This characteristic trait indicates that E484K coexisting with N501Y also serves to increase the transmissibility of SARS-CoV-2. Our results suggest that, like D614G, E484K achieves this by promoting the S conformation to shift from predominantly “all-RBD-down” hACE2-inaccessible closed state to “one/two-RBD-up” hACE2-accessible partially open states, thus exposing the RBD and facilitating receptor binding of S. Similar to that of D614G, emerged E484K carrying S variants likely promote transmissibility by adopting more dynamics-decelerated or implied stability-enhanced fusion-promoting hACE2-accessible conformations, revealed by our smFRET results. Notably, E484K was also identified as an escape mutation, as it enables the virus to escape host antibody recognition ^9,24,68-70^. This immune-evasive trait could be caused by the reversal of electric charge at position 484 due to the E-to-K mutation, which creates a disadvantaged electrostatic interaction site that likely weaken the binding efficiency of anti-RBD antibodies. The shift of electric charge could potentially cause local conformational changes in the RBD ^16^, which exceeds the capacity of our current smFRET setup. It is possible that this unique gain-of-function mutation was selected to reconcile the cost of exposing the RBD, which is a primary target of highly potent neutralizing antibodies. It could explain why the original Alpha variant, identified in the UK, did not contain the E484K but has since adopted it. By the time of performing this study, SARS-CoV-2 has modified its survival strategy by adopting E484K in addition to D614G to confer higher transmissibility (likely complemented with N501Y) and protect S against antibodies. As the delta variant currently out-compete others as the dominant pandemic form, SARS-CoV-2 could have shifted its survival strategy again, which requires further investigation beyond the scope of this study.

In conclusion, we used smFRET to reveal real-time conformational populations adopted by virus-associated S variants and experimentally identified conformation-interconverting temporal dynamics. We found that S evolves in favor of the hACE2-binding competent “RBD-up” conformations, and “RBD-up” conformations exhibit decelerated dynamics than “RBD-down.” Our results explain the inconsistent results regarding the hACE2-binding ability of S variants and S stability. Our findings improve the molecular understanding of earlier S variants underlying immune evasion and virus transmissibility and further provide the molecular basis of our knowledge and prediction on SARS-CoV-2 evolution, informing S-centric interventions to curtail the COVID-19 pandemic.

## ACKNOWLEDGMENTS

We thank Dr. David Derse for sharing HIV In-GLuc plasmids. We thank Dr. Scott Blanchard for sharing smFRET software and guiding the design of the prism-TIRF microscope. This work was supported by The University of Texas Health Science Center at Tyler to M.L., and by NIH/NIAID grant R01 AI163395 to W.M. This work was partially supported by le Ministère de l’Économie et de l’Innovation du Québec, Programme de soutien aux organismes de recherche et d’innovation to A.F. and by the Sentinelle COVID Quebec network led by the LSPQ in collaboration with Fonds de Recherche du Québec Santé (FRQS) to A.F. A.F. is the recipient of Canada Research Chair on Retroviral Entry no. RCHS0235 950-232424.

## AUTHOR CONTRIBUTIONS

M.L. and W.M. designed the studies. Z.Y. and Y.H performed mutagenesis. Z.Y., Y.H, and M.L. designed and performed virus infectivity assays. M.L., Z.Y. and Y.H generated fluorescently labeled viruses, performed smFRET data acquisition and analyzed the data. M.L., Z.Y. and Y.H made figures, tables, and models. S.D. and A.F. provided hACE2. M.L., Z.Y. and W.M. wrote the manuscript with inputs from other authors.

## METHODS

No statistical methods were used to predetermine sample size. The experiments were not randomized and investigators were not blinded to allocation during experiments and outcome assessment.

### Cell Lines

293T (or HEK293T) cells, 293T-ACE2, and Expi293F cells were cultured in DMEM media, supplemented with 10% FBS, 100 U/ml penicillin/ streptomycin, 2 mM L-glutamine, and in the presence of 5% CO_2_. Cell culture media was exchanged with fresh media before transfection. 293 cells, human embryonic kidney cells, were obtained from ATCC (cat # CRL-1573™). 293T cells were 293 cells derivative with the simian virus 40 T-antigen inserted. 293T-ACE2 cells, derived from 293T, stably express human ACE2 ^71^. Expi293F cells, derived from the 293 cells, were purchased from ThermoFisher Scientific (cat # A14528; RRID: CVCL_D615).

### Construction of full-length untagged and tagged SARS-CoV-2 spike (S) variants

Our current FRET-engineered system uses enzymatic labeling that introduces peptide tags located sterically further from any of accumulated mutations on S variants. Untagged and double-tagged SARS-CoV-2 spike variants were generated based on a template full-length pCMV3-SARS-CoV-2 Spike (codon-optimized, Sino Biological, cat # VG40589-UT) plasmid that has translated amino acid sequence identical to QHD43416.1 (GenBank). As described previously ^55^, peptide-based labeling tags Q3 (GQQQLG) and A4 (DSLDMLEM) ^59,60^ were engineered into the parental S D614 (S_D614_, first identified in Wuhan) and tested variants in this study at positions before and after the receptor-binding motif (RBM) to avoid interfering with the binding of cellular receptor human ACE (hACE2) (Figure 1C). Insertion of a pair of 427-Q3/556-A4 into S did not compromise S-dependent lentivirus infectivity and this double-tagged FRET-engineered S_D614_ (designated as S_D614_ Q3/A4) were constructed for smFRET imaging of S_D614 ^55^_. D614G point mutation was introduced into both untagged full-length pCMV3 S_D614_ and double-tagged S_D614_ Q3/A4 constructs to generate both untagged and tagged S_G614_ variants. E484K and N501Y were further introduced into the untagged S_G614_ individually and together. In the case of S Alpha (pcDNA3.1 S_Alpha_, codon-optimized) variant, DNA sequences encoding untagged or double-tagged (with tags positioning at the exact positions as in tagged S_D614_ or S_G614_) were synthesized by GeneScript and cloned into the pcDNA3.1 vectors. E484K point mutation was introduced into both untagged and tagged pcDNA3.1 S_Alpha_ by site-specific mutagenesis.

### Viral infectivity measurements

The infectivity of lentivirus particles carrying S proteins (including variants) on the surface was evaluated using a vector containing an HIV-1 long terminal repeat (LTR) that expresses a Gaussia luciferase reporter (HIV-1-inGluc) ^72,73^. 293T cells were transfected at 60–80 % confluency with the plasmid encoding indicated full-length SARS-CoV-2 S glycoproteins, the plasmid encoding an intron-regulated Gluc (HIV-1-inGluc), and a plasmid pCMV delta R8.2 encoding HIV-1 GagPol (Addgene, plasmid # 12263) using FuGENE 6 (Promega, # E2311). 40 hours post-transfection, lentivirus particles were harvested, filtered through a 0.45 mM filter (Pall Corporation), and titered on target cells 293T-ACE2 that endogenously express hACE2. Lentivirus infectivity was determined 48 hours post-infection by measuring Gaussia luciferase activity in the 293T-ACE2 supernatant using a Pierce™ Gaussia luciferase flash assay kits (ThermoFisher Scientific, cat# 16158).

### hACE2 expression and purification

The human ACE2 (hACE2) expression construct was synthesized (Gene Universal Inc, Newark DE) and subcloned into corresponding pVRC8400 vectors. Monomeric (residues 1-615) human ACE2 proteins were produced as described previously ^74^. Briefly, DNA sequence encoding monomeric hACE2 followed by an HRV3C cleavage site, monomeric Fc tag, and 8xHisTag at the 3’-end were synthesized and cloned into the pVRC8400 vector. The proteins were expressed by transiently transfecting Expi293F cells and then purified by Protein A affinity columns. The Fc and 8xHis tags were removed by overnight HRV3C digestion at 4 °C. hACE2 proteins were further purified over a Superdex 200 16/60 column in the pH7.5 buffer solution containing 5 mM HEPES and 150 mM NaCl.

### Preparation of lentivirus particles carrying S proteins for smFRET

Lentivirus particles carrying S proteins (the ancestor and variants) on the surface were prepared similarly as previously described for the parental S_D614 ^55^_ and HIV-1 ^57,58,75^. Briefly, lentivirus particles were produced by pseudo-typing an HIV-1 core with full-length S proteins. To conduct studies of S variants by smFRET, we introduced the labeling peptide tags Q3 and A4 at the same positions (427 and 556, respectively) in S1 of S_G614_, S_Alpha_, E484K carrying S_Alpha+E484K_ variants as in previously reported S_D614_^55^. Lentivirus particles used for smFRET imaging were made in 293T cells by transfecting a 20-fold excess of plasmid encoding S_D614_, S_G614_, S_Alpha_, or S_Alpha+E484K_ over their corresponding 427-Q3/556-A4 to ensure that, on average, every virus particle carries a single Q3/A4-tagged S protomer. Plasmids encoding indicated untagged wildtype S (or variants), the corresponding double-tagged/FRET-engineered S, and a lentivirus packaging delta R8.2 were used at a ratio of 20:1:21, respectively. 293T cells were transfected using FuGENE 6 with above-indicated plasmids to express spikes on the HIV-1 lentivirus particles. Particles were harvested 40 hours post-transfection, filtered through a syringe filter with 0.45 μm pore size, and sedimented through a 15 % sucrose cushion at 25,000 rpm for 2 hours. The virus pellets were then re-suspended in pH 7.5 50 mM HEPES containing 10 mM MgCl_2_ and 10 mM CaCl_2_. Of note, the strategy of using an excess of plasmid-encoding wildtype vs. trace amounts of a plasmid expressing labeling-tag-carrying S ensures the production of virus particles that contain, on average, only a single FRET-engineered S protomer. While most virus particles carry untagged S, among the small portion of virus particles containing tagged S, more than 95 % will carry one dually tagged protomer while the other two protomers remain wildtype. The same strategy has been used in our previous smFRET studies ^55,57,58,75^.

### Fluorescently labeling S proteins on lentivirus particles

Lentivirus particles were fluorescently labeled through site-specific labeling based on enzymatic reactions, as previously described ^55,57,58,75^. Briefly, transglutaminase transferred a Cy3B(3S) derivative (LD555-cadaverine) from the cadaverine conjugate to the central glutamine residue of the Q3 (GQ<u>Q</u>QLG) tag in S1. The AcpS enzyme catalyzed the attachment of the Cy5 derivative (LD655-CoA) to the serine residue of the A4 tag (D<u>S</u>LDMLEM). LD555-cadaverine (0.5 μM, Lumidyne Technologies), LD655-CoA (0.5 μM, Lumidyne Technologies), transglutaminase (0.65 μM, Sigma Aldrich), and AcpS (5 μM, home-made) were added to the above lentivirus particles prepared for smFRET, and incubated at room temperature overnight. LD555-cadaverine and LD655-CoA were synthesized by Scott Blanchard laboratory (Lumidyne Technologies) based on new generation photostable Cy3/Cy5 organic dyes ^76^. DSPE-PEG2000-biotin (0.02 mg/ml, Avanti Polar Lipids) was then added to the reaction mix and incubated for 30 min at room temperature with rotation. Labeled lentivirus particles were then purified to eliminate excess free dyes and lipids by ultracentrifugation for one hour at 40,000 rpm over a 6%-18% Optiprep (Sigma Aldrich) gradient. Purified fluorescently-labeled lentivirus particles were stored at -80 °C.

### smFRET imaging data acquisition

All smFRET imaging experiments were performed on a home-built prism-based total internal reflection fluorescence (TIRF) microscope. Lentiviruses particles carrying fluorescently labeled spike proteins were immobilized on PEG-passivated quartz slides coated with streptavidin. The evanescent field was generated at the interface between the quartz slide and the virus-containing sample solution by prism-based total internal reflection, with 532-nm CW laser excitation (Laser Quantum). The donor fluorophore (LD555) labeled on the virus is excited by the generated evanescent field and can transfer energy at its excited states to the neighboring acceptor fluorophore (LD655). Fluorescence from both fluorophores was collected through a 1.27-NA 60 x water-immersion objective (Nikon) and then optically separated by a 650 DCXR dichroic filter (Chroma) mounted on a MultiCam LS image splitter (Cairn Research). Fluorescence of donor (ET590/50, Chroma) and acceptor (ET690/50, Chroma) was separately and simultaneously recorded on two synchronized ORCA-Flash4.0 V3 sCMOS cameras (Hamamatsu) at 25 Hz for 80 seconds. Fluorescently labeled virus samples were imaged in pH 7.4, 50 mM Tris buffer containing 50 mM NaCl, a cocktail of triplet-state quenchers, and an oxygen-scavenger system^77^. Where indicated, the conformational effects of hACE2 on S proteins were conducted by pre-incubating fluorescently labeled viruses with 200 μg/ml hACE2 for 90 mins at room temperature before imaging and smFRET imaging data were taken in the continued presence of 200 μg/ml hACE2.

### smFRET quantification and statistical analysis

The analysis of smFRET data was performed using a MATLAB-based customized SPARTAN software package ^78^. Both donor and acceptor fluorescence intensity of each labeled virus were recorded over 80 seconds. The background signals at the single-molecule level were first identified based on the single-step fluorophore bleaching point and subtracted from the recorded signals. For each recorded virus, donor and acceptor fluorescence intensity traces (fluorescence time trajectory) were then extracted and corrected for the donor to acceptor crosstalk. The energy transfer efficiency from the donor to the acceptor (FRET values, simplified as FRET in graphs) was calculated over time according to FRET= I_A_/(γI_D_+I_A_), where I_D_ and I_A_ are the fluorescence intensities of donor and acceptor, respectively, and γ is the coefficient correcting for the difference in detection efficiencies of donor and acceptor. FRET traces (FRET values as a function of real-time) monitor the relative donor-to-acceptor distance over time, which reflect conformational dynamics of the donor/acceptor-labeled spike on the lentivirus in real-time. Virus particles that fall into any of the following categories were automatically excluded from our data analysis: 1) lacking either donor or acceptor or both fluorophores; 2) containing multiple labeled protomers; 3) containing more than one labeled spike on a single virus. In addition, FRET traces from individual viruses that meet our initial selection criteria and display sufficient signal-to-noise (S/N) ratio and anti-correlated fluctuations in donor and acceptor fluorescence were manually picked for data analysis. The anti-correlated feature directly defines FRET-indicated conformational states, which indicate fully active/dynamic spikes on virus particles. Thus, only one FRET-labeled protomer within one S trimer on one lentivirus particle (1-protomer:1-trimer:1-virus particle) showing clear anti-correlated features of donor and acceptor fluorescence and single fluorescence bleaching point passed our filters. FRET traces included in our analysis were then compiled into FRET histograms, which reflect conformational ensembles. Based on visual inspection of fluorescence and FRET traces that revealed direct observations of state-to-state (conformation-to-conformation) transitions, FRET histograms were fitted into the sum of four Gaussian distributions using the least-squares fitting algorithm in MATLAB. Each gaussian represents one FRET-defined conformational state of S proteins, and the area under each Gaussian curve estimates the relative probability of S occupying each conformation. The relative occupancy of each conformational state was used to evaluate the difference in relative free energies between states *i* and *j*, according to *ΔG°*_*ij*_*=-k*_*B*_*Tln(P*_*i*_*/P*_*j*_), where *P*_*i*_ and *P*_*j*_ are the occupancy of *i*th and *j*th states, respectively, *k*_*B*_ is the Boltzmann constant, and *T* is the temperature in kelvin. FRET traces compiled in the FRET histogram under indicated experimental conditions were idealized using a segmental *K*-means algorithm ^63^ with a 4-state Hidden Markov Model – providing the simplest explanation of our data. The frequency and order of idealized state-to-state transitions were displayed in a transition density plot (TDP). Dwell times of S in each states were compiled into survival probability plots which were fitted to the sum of two exponential distributions (*y(t)* = *A*_1_ exp ^−*k*1*t*^ + *A*_2_ exp ^−*k*2*t*^), where *y(t)* is the probability and *t* is the dwell time. The transition rates were determined by averaging the two rate constants weighted by their amplitudes.

## FIGURES AND LEGENDS

**Figure S1.**
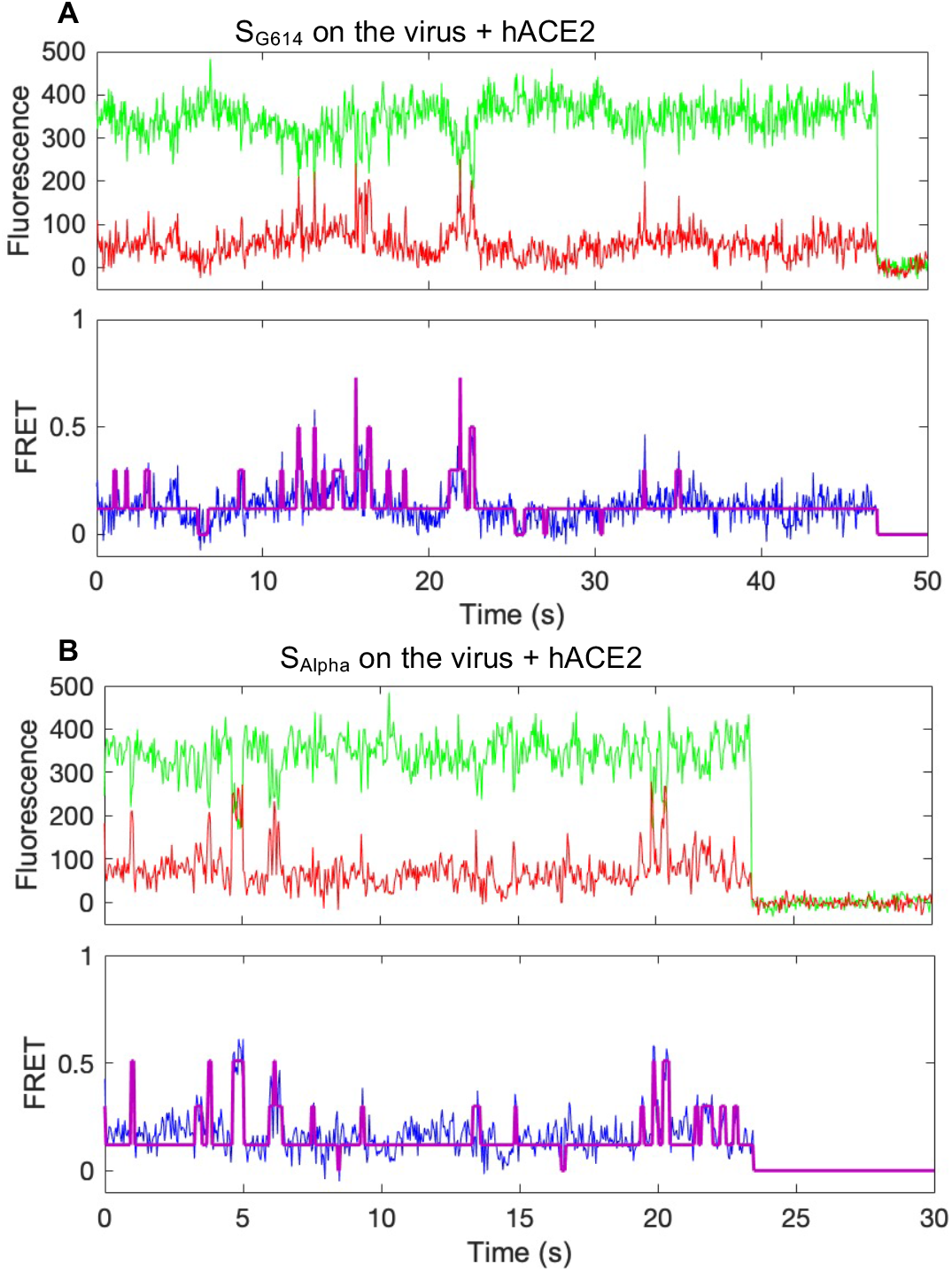
Receptor hACE2 activates spike variants S_G614_ and S_Alpha_ towards fusion-competent “all-RBD-up” conformation. (Related to Figures 2 and 3) (**A, B**) Example fluorescence traces (LD555, green; LD655, red) and resulting quantified low-FRET dominated traces (FRET efficiency, blue; hidden Markov model initialization, red) of lentivirus-embedded spikes variants G614 (**A**) and Alpha (**B**) in response to hACE2.

**Figure S2.**
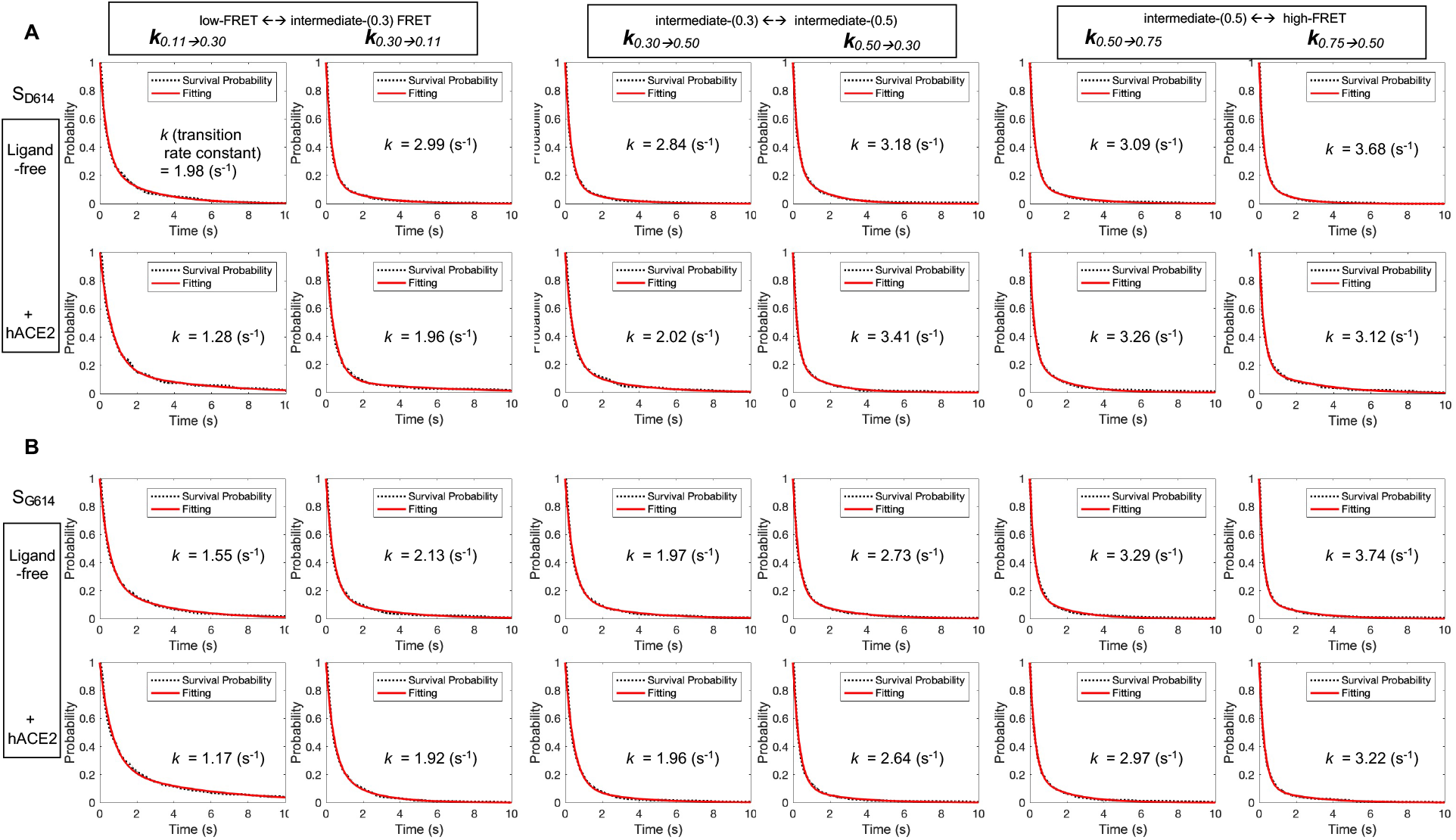
Kinetic analysis of spike proteins S_D614_ and S_G614_ on lentivirus particles and modulation by hACE2. (Related to Figure 4). (**A, B**) Survival probability plots of four FRET-indicated conformations of lentivirus particles-associated spike proteins S_D614_ (**A**) and S_G614_ (**B**) in the absence (ligand-free, top row) and the presence of 200 μg/ml hACE2 (bottom row). The survival probability plot is the time-duration distribution of spikes dwelling on one conformation before directionally transition to the other. Transition rates (summarized in Figure 4B) were estimated by double exponential-fitting of survival probability plots.

**Figure S3.**
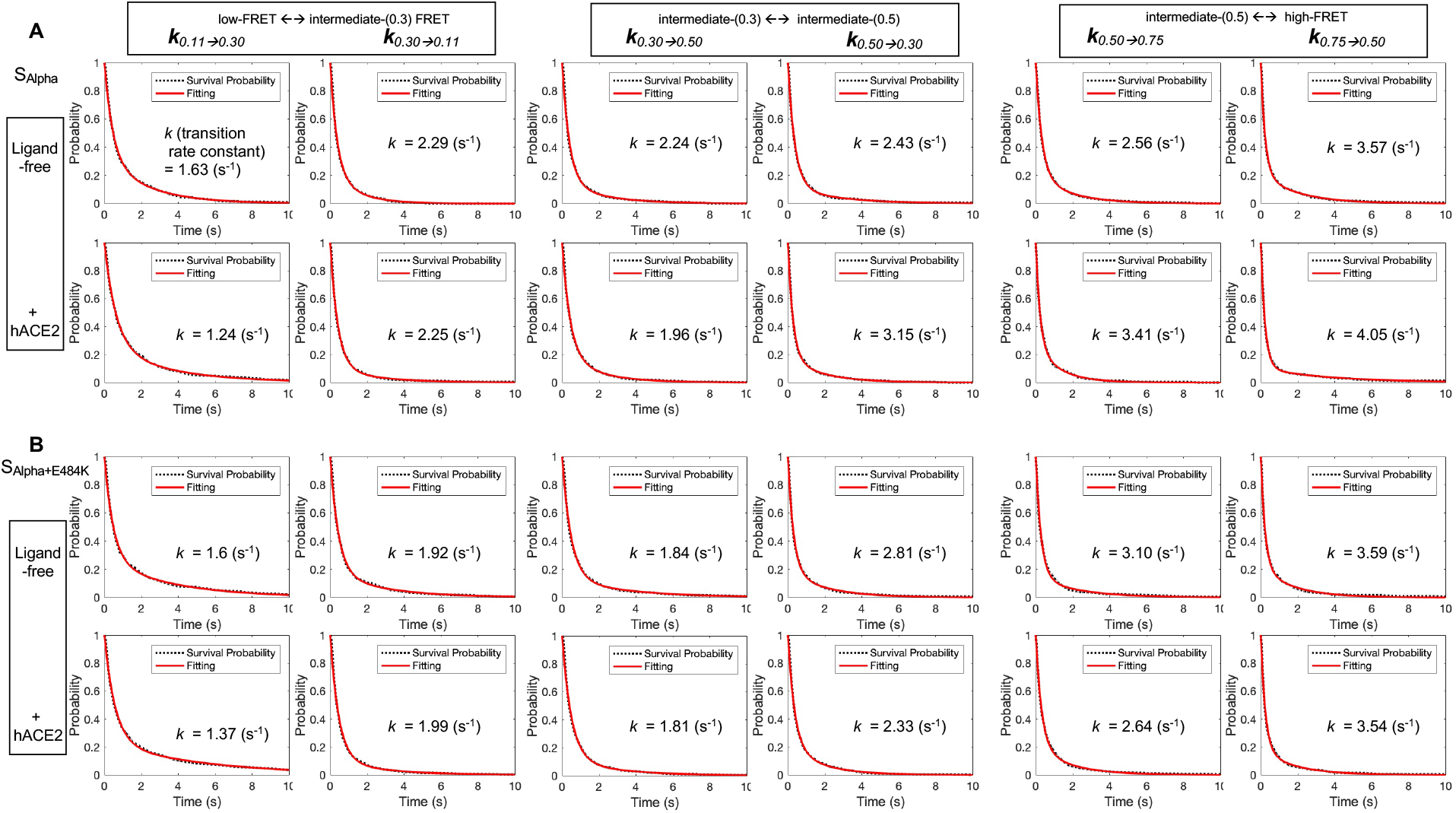
Kinetic analysis of spike variants S_Alpha_ and S_Alpha+E484K_ on lentivirus particles, and modulation by hACE2. (Related to Figure 4). (**A, B**) Survival probability plots as in Figure S2 for spike variants S_Alpha_ (**A**) and S_Alpha+E484K_ (**B**) give rise to the estimation of transition rates among different conformations of spike variants. Transition rates were summarized in Figure 4B.

**Table S1.** **Relative probability and determining parameters of four FRET-defined conformational states occupied by spike variants on lentivirus particles. (Related to Figures 2 - 5)**. Conformational distribution-indicated FRET efficiency histograms were fitted into the sum of four distinct Gaussian distributions (μ, mean or expectation; σ, standard deviation) for each conformation. Parameters (μ and σ) were based upon the observation of original FRET efficiency traces and were further determined using hidden Markov modeling. The probability of conformations that spikes occupy was presented as mean ± s.e.m., determined from three independent measurements. R-squared values were evaluated to indicate the goodness of fit.

| Spike variants on lentivirus particles | Curve fitting R-squared | low-FRET | intermediate-(0.3) FRET | intermediate-(0.5) FRET | high-FRET |
| --- | --- | --- | --- | --- | --- |
| | | $\mu$: 0.11+/-0.01 | $\mu$: 0.30+/-0.01 | $\mu$: 0.50+/-0.02 | $\mu$: 0.75+/-0.03 |
| | | $\sigma$: 0.08+/-0.01 | $\sigma$: 0.08+/-0.01 | $\sigma$: 0.10+/-0.01 | $\sigma$: 0.10+/-0.01 |
| $S_{D614}$ | | | | | |
| Ligand-free | 0.9941 | 25% +/- 7% | 25% +/- 11% | 38% +/- 13% | 12% +/- 7% |
| + hACE2 | 0.9928 | 42% +/- 10% | 27% +/- 11% | 21% +/- 12% | 10% +/- 3% |
| $S_{G614}$ | | | | | |
| Ligand-free | 0.9967 | 26% +/- 8% | 34% +/- 13% | 29% +/- 12% | 11% +/- 6% |
| + hACE2 | 0.9963 | 44% +/- 4% | 20% +/- 6% | 25% +/- 7% | 11% +/- 3% |
| $S_{Alpha}$ | | | | | |
| Ligand-free | 0.9890 | 19% +/- 10% | 29% +/- 14% | 34% +/- 15% | 18% +/- 10% |
| + hACE2 | 0.9956 | 41% +/- 10% | 26% +/- 10% | 24% +/- 10% | 9% +/- 5% |
| $S_{Alpha+E484K}$ | | | | | |
| Ligand-free | 0.9890 | 23% +/- 8% | 38% +/- 11% | 29% +/- 11% | 10% +/- 5% |
| + hACE2 | 0.9956 | 49% +/- 9% | 22% +/- 13% | 22% +/- 13% | 7% +/- 5% |

